# Systematic Benchmarking of Kinase Bioactivity Models Across Splitting Strategies and Protein Representations

**DOI:** 10.64898/2026.04.20.719590

**Authors:** Joshua M. Abbott

## Abstract

Machine learning models for protein–ligand bioactivity prediction are increasingly used in computational drug discovery. However, reported benchmark performance is often sensitive to evaluation design. To further understand evaluation design strategies, we present a systematic evaluation of seven machine learning architectures for kinase inhibitor bioactivity prediction, spanning classical baselines (Random Forest, XGBoost, ElasticNet, multi-layer perceptron) and advanced neural approaches (Graph Isomorphism Network, ESM-2 protein embedding MLP, and a GNN-ESM fusion model). Using a curated ChEMBL-derived kinase activity dataset of 352,874 records across 507 human protein kinase targets, we evaluated all models under three splitting strategies of increasing stringency: random, scaffold-based (Bemis–Murcko), and target-held-out. We observed that Random Forest with Morgan fingerprints achieves near-equivalent or superior performance to all neural architectures under scaffold and target-based evaluation. On target-held-out splits frozen ESM-2 embeddings showed worse generalization, with ESM-FP MLP exhibiting the largest performance degradation. Learned graph representations (GIN) do not outperform fixed 2048-bit ECFP4 fingerprints at this data scale, and tree-based uncertainty methods outperform MC-Dropout implementations tested here on calibration and selective prediction metrics. A JAK kinase subfamily case study shows that protein-aware models achieved 79% top-1 selectivity accuracy versus 52% for pooled fingerprint models. However, stronger baselines using explicit target identity achieved 83–84%, indicating that ESM-2 embeddings in this study functioned primarily as an implicit target identifier. These results indicate that evaluation methodology and statistical rigor are major determinants of reported performance in bioactivity prediction.

**Benchmark design overview:** 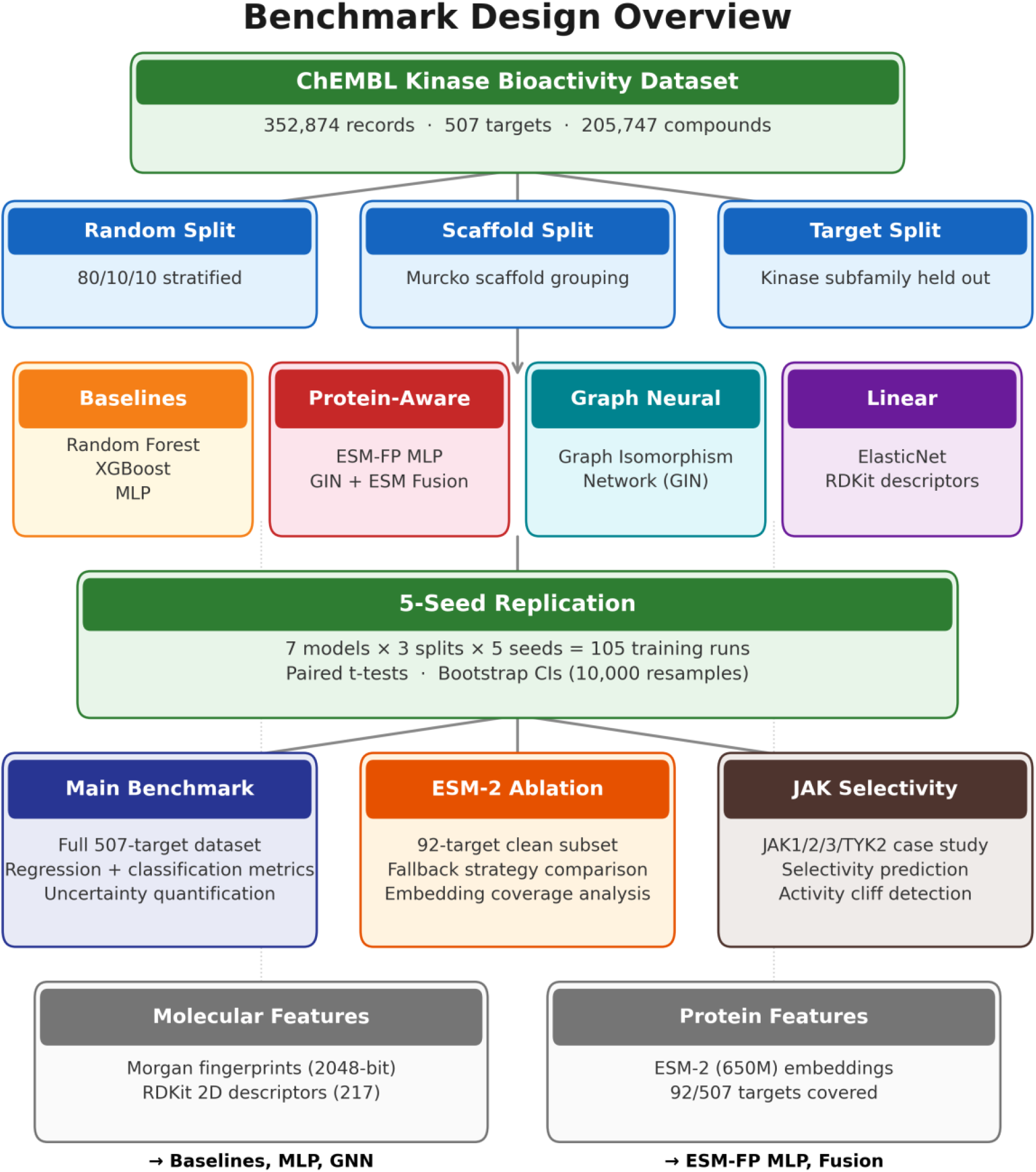

A curated ChEMBL kinase bioactivity dataset (352,874 records, 507 targets) was evaluated under three splitting strategies of increasing stringency. Seven model architectures spanning baselines, protein-aware, and graph neural approaches were each trained under 5-seed replication (105 total runs), with results analyzed across three complementary branches: the main 507-target benchmark, ESM-2 embedding ablation studies on a clean 92-target subset, and a JAK-family selectivity case study with stronger target-conditioned baselines

## Introduction

Protein kinases constitute one of the most therapeutically important target families in drug discovery, with over 70 FDA-approved kinase inhibitors as of 2024, primarily for oncology and inflammatory disease indications^1,2^. The conserved ATP-binding architecture of kinases makes selectivity optimization a central challenge in kinase drug discovery, as inhibitors frequently exhibit promiscuous binding across the family. Predicting bioactivity of small molecules against kinase targets is therefore an important problem for computational drug discovery.

Machine learning approaches for molecular property prediction have made substantial advances in recent years, with published methods spanning traditional cheminformatics descriptors^3,4^, learned molecular graph representations^5,6^, and protein-aware architectures that incorporate target sequence information through protein language models^7,8^. Recent evidence suggests that pLM embedding quality is heterogeneous across the proteome, with a substantial fraction of proteins receiving low-confidence representations that may degrade downstream predictions^9^. Beyond representation quality, however, the evaluation methodology employed in many published benchmarks raises concerns about the generalizability of reported results. Random train/test splits, which remain the standard in many studies, often overestimate prospective performance because close structural analogs and repeated target identities appear across partitions, inflating performance metrics substantially^10–12^.

Scaffold-based splitting, which groups compounds by Bemis–Murcko scaffolds and assigns entire scaffold clusters to a single partition, provides a more realistic estimate of model performance on novel chemical matter^13^. Target-based splitting, which holds out entire protein subfamilies, tests the most challenging form of generalization, prediction for targets the model has not seen during training. Despite their importance, systematic comparisons across all three splitting strategies with matched datasets and evaluation protocols remain uncommon in the literature.

Several studies have demonstrated that simple baselines can be competitive with deep learning architectures when evaluated rigorously. Work by Mayr and colleagues conducted a large-scale comparison on ChEMBL and found that while deep learning methods significantly outperformed classical approaches on some tasks, the advantage was task-dependent^14^. Yang et al. showed that their directed message-passing neural network (D-MPNN) outperformed fingerprint baselines on MoleculeNet benchmarks, but the magnitude of improvement varied considerably across datasets and splitting strategies^6,15^. Luo et al. demonstrated that current deep learning models for kinase inhibitor affinity prediction show essentially no generalization when information leakage is eliminated. The conditions under which more complex architectures outperform fingerprint-based baselines in kinase bioactivity prediction remain incompletely resolved^16^.

Uncertainty quantification (UQ) is equally important for practical deployment. A model that provides well-calibrated confidence estimates enables medicinal chemists to prioritize high-confidence predictions for experimental validation and flag uncertain predictions for additional computational analysis^17,18^. The relative performance of different UQ approaches (tree variance, quantile regression, deep ensembles, MC-Dropout^19^) across model types has not been systematically benchmarked in the context of kinase affinity prediction.

In this work, we seek to understand these gaps through a systematic evaluation of seven machine learning architectures for kinase inhibitor bioactivity prediction, using a curated dataset of 352,874 bioactivity records from ChEMBL spanning 507 human protein kinase targets^20^. We initially evaluated seven models. However, because ElasticNet performed substantially worse than the remaining six across all splits (scaffold RMSE = 1.274 vs. next-worst 0.961), it is reported in supporting data and omitted from the main comparison tables for readability. We evaluate all models under random, scaffold, and target-held-out splits with consistent metrics, bootstrap confidence intervals, uncertainty quantification, and error analysis. A case study on the JAK kinase family (JAK1, JAK2, JAK3, TYK2) provides compound-level insight into model failure modes and reveals a specific niche where protein-aware models appear to provide their clearest advantage, selectivity prediction across closely related targets. In this paper, we quantified how strongly reported performance depends on splitting strategy across all model types, with 5-seed replication revealing a false positive in single-seed comparisons. We performed a systematic comparison of seven architectures under three splitting strategies with 5-seed replication, an evaluation of frozen ESM-2 protein embeddings with ablation studies addressing incomplete coverage, uncertainty quantification benchmarking across model families, and a JAK-family selectivity case study with stronger target-conditioned baselines.

## Methods

### Dataset Construction

Bioactivity data were retrieved from ChEMBL 36 for all human protein kinase targets identified via Gene Ontology molecular function annotations (GO:0016301, GO:0004672) and name-based filtering, yielding 507 unique single-protein kinase targets^20^. Protein-aware models additionally required curated UniProt sequences for ESM-2 embedding computation, sequences were successfully retrieved for 92 of the 507 targets. All models were trained and evaluated on the full 507-target dataset. For the 415 targets without sequences, protein-aware models received a shared fallback embedding corresponding to an arbitrary kinase. Activity measurements (IC_50_, K_i_, K_D_) were filtered to exact values only from high-confidence assays (confidence score ≥ 7) with pChEMBL values available.

Molecules were standardized using RDKit: salt removal, charge neutralization, canonical SMILES generation, and molecular weight filtering (100–900 Da, maximum 100 heavy atoms)^4^. Activity values were converted to a unified pActivity scale (pActivity = −log_10_ M). For duplicate compound-target-assay type triples, the median pActivity was taken. Compounds with ≥ 3 replicate measurements and standard deviation > 1.0 pActivity units were flagged in metadata as potentially noisy but retained in all analyses (1,965 measurements, 0.56% of data). The final curated dataset comprises 352,874 records spanning 205,747 unique compounds, pActivity range 3.0–11.0 (mean 7.0, SD 1.28). The dataset is enriched for active compounds (77.3% with pActivity ≥ 6.0).

### Splitting Strategies

The random split (80/10/10 train/validation/test) is stratified by target to maintain representation across partitions. The scaffold split groups compounds by genericized Bemis–Murcko scaffolds^13^ were computed via RDKit. Entire scaffold groups are assigned to a single partition via greedy assignment sorted by group size, preventing scaffold leakage. The target split holds out entire kinase subfamilies defined using ChEMBL protein family annotation for testing. The exact train/validation/test family assignments are provided in the code repository. Split sizes are: random (282,299/35,287/35,288), scaffold (282,301/35,288/35,285), and target (251,532/65,811/35,531). The canonical benchmark uses seed 42; the single-partition design is noted as a limitation.

### Feature Representations

#### Morgan Fingerprints (ECFP4)

Computed using RDKit’s rdFingerprintGenerator.GetMorganGenerator API^3,4^ with default parameters (includeChirality = False). Used as input for Random Forest, XGBoost, and MLP models.

#### RDKit 2D Descriptors

217 physicochemical and topological descriptors computed using RDKit’s CalcMolDescriptors^4^. Descriptors with > 5% missing values were removed, remaining NaNs imputed with column medians. Used as input for ElasticNet.

#### Molecular Graphs

Atom-level features 35 dimensions: element type, degree, formal charge, hybridization, aromaticity, chirality and bond-level features bond type, stereochemistry, conjugation, ring membership. Used for GIN and Fusion models.

#### Protein Embeddings

Protein target sequences were retrieved from UniProt via ChEMBL cross-references for 92 of the 507 kinase targets (18.1%). These sequences were embedded using frozen ESM-2 (esm2_t33_650M_UR50D, 650M parameters)^7,8^, truncated to 1,022 residues (N-terminal; affecting 19 of 92 sequences) and mean-pooled over residue positions to yield 1,280-dimensional target embeddings. For the remaining 415 targets, a shared fallback embedding was used, row 0 of the embedding matrix, corresponding to an arbitrary kinase. Used for ESM-FP MLP and Fusion models.

### Model Architectures

#### Baseline Models

##### Random Forest

500 trees, no depth limit, min_samples_leaf = 2, max_features = sqrt. Uncertainty via tree prediction variance^22^.

##### XGBoost

500 trees, max_depth = 6, learning_rate = 0.1, subsample = 0.8^23^. Uncertainty via quantile regression (α = 0.05, 0.95).

##### ElasticNet

L1/L2 regularized linear regression on standardized RDKit descriptors. Uncertainty via bootstrap resampling (100 models). Hyperparameters were selected via validation-set optimization (α = 0.001, l1_ratio = 0.1).

##### MLP

Two hidden layers [256, 128], ReLU, Adam optimizer, batch_size = 512, early stopping (patience = 10). Uncertainty via ensemble of 3 independently initialized models.

#### Advanced Neural Models

##### ESM-FP MLP

Concatenates Morgan fingerprints (2,048-dim) with ESM-2 embeddings (1,280-dim) for 3,328-dim input. Architecture: [Linear → ReLU → Dropout → BatchNorm] × 2 → Linear → ReLU → Dropout → Linear(1). Uncertainty via MC-Dropout (10 forward passes)^19^.

##### GIN (Graph Isomorphism Network)

3 GINConv layers with 128-dim hidden states, global mean + max pooling (256-dim), followed by MLP prediction head^5^. Uncertainty via MC-Dropout.

##### GIN + ESM-2 Fusion

Dual-branch: GIN ligand branch (→ 256-dim) concatenated with ESM-2 protein branch (1,280 → 256-dim), fed through shared MLP head. Uncertainty via MC-Dropout.

All deep models use AdamW optimizer with cosine annealing learning rate schedule and early stopping (patience = 15 epochs on validation RMSE). Training was performed on AWS EC2 GPU instances. To make comparisons as fair as possible, all model classes were tuned against the same validation split using model-specific search spaces. Systematic hyperparameter searches were conducted via grid search for XGBoost with 20 configurations over tree depth, learning rate, and number of estimators and for ElasticNet 35 configurations over regularization strength and L1 ratio. Random Forest, MLP, and deep models used fixed architectures with early stopping as implicit regularization. Full search spaces, selected values, and sensitivity analyses are provided in Table S2.

### Evaluation Metrics

Regression: RMSE, MAE, R^2^, Pearson r, Spearman ρ. Classification (pActivity ≥ 6.0 threshold): AUPRC and Precision at K were used as primary classification metrics, with AUROC included for completeness. Uncertainty: miscalibration area, error-uncertainty, Spearman correlation, selective prediction improvement at 50% retention. All metrics on held-out test sets.

### Statistical Framework

To differentiate between training stochasticity from test-set sampling variability, we employ two complementary approaches. Multi-seed training (5 seeds: 42, 123, 456, 789, 1024): each model is trained 5 times on each split, yielding 105 experiments. We report mean ± SD across seeds and use paired t-tests (df = 4) for pairwise comparisons, with effect sizes reported as absolute ΔRMSE and percent difference. Five seeds were chosen as a practical compromise between computational cost (105 training runs totaling ∼160 GPU-hours) and statistical stability. Per-seed results (Appendix A) confirm that cross-seed variance is well-characterized at this sample size. Partition stochasticity is not addressed by multi-seed replication and remains a limitation. Bootstrap confidence intervals (10,000 resamples) for each experiment, we resample test predictions to estimate CIs and run paired bootstrap tests. Given large test sets, bootstrap CIs are narrow (±0.008 RMSE typical). Where multi-seed and bootstrap results diverge, we defer to multi-seed results. For nonparametric sensitivity check, we examined Wilcoxon signed-rank tests on the 5-seed comparisons for n = 5, the minimum achievable two-tailed p-value is 0.0625. All comparisons where all 5 seed-level differences favored one model achieved W = 0, p = 0.0625.

### JAK Case Study Design

The Janus kinase (JAK) family, JAK1, JAK2, JAK3, and TYK2, was selected for a compound-level case study because it offers the richest analytical opportunity in the dataset: 36,059 records across 4 well-characterized members, with 2,498 compounds tested against all 4 JAKs. Multiple FDA-approved JAK inhibitors like tofacitinib, baricitinib, ruxolitinib provide clinical context. The high compound overlap between members enables a selectivity prediction analysis that tests whether protein-aware models can distinguish which JAK member a compound preferentially inhibits.

Activity cliff pairs were identified as structurally similar compounds (Tanimoto ≥ 0.85) with large activity differences (|ΔpActivity| ≥ 1.5 log units). Selectivity prediction was evaluated by asking each model to rank JAK family members by predicted potency for each multi-JAK compound; top-1 accuracy measures the fraction where the model correctly identifies the most potently inhibited JAK member. All models are pooled multi-target regressors trained on compound, target, pActivity triples across all kinases, not separate per-target models. Ligand-only models (RF, XGBoost, MLP) receive identical Morgan fingerprint input for a given compound regardless of the target, and produce near-identical predictions across JAK members, their ∼52% top-1 accuracy reflects this architectural limitation and serves as a baseline for the selectivity task. Protein-aware models (ESM-FP MLP, Fusion) receive different input for different targets because the ESM-2 embedding changes, enabling cross-target discrimination.

## Results

### Regression Performance Across Splitting Strategies

**Table 1** presents regression metrics for the six primary comparison models across three splitting strategies (**Figure 1**). Deep model values are reported as multi-seed means ± SD (5 seeds); tree-based models show near-zero cross-seed variance (±0.002). On random splits, ESM-FP MLP achieves the best RMSE (0.777 ± 0.002), outperforming Random Forest by ΔRMSE = 0.042 (5.1%, paired t-test p < 0.001) and MLP by ΔRMSE = 0.049 (6.0%, p < 0.001). However, model rankings change substantially under more stringent evaluation. On scaffold splits, MLP (0.901 ± 0.003) and ESM-FP MLP (0.902 ± 0.004) are not statistically different (ΔRMSE = −0.001, p = 0.575) correcting a single-seed bootstrap analysis that incorrectly suggested a significant 0.9% ESM-FP MLP advantage (p = 0.003). On target splits, Random Forest (1.066 ± 0.002) significantly outperforms ESM-FP MLP by ΔRMSE = 0.113 (10.6%; p < 0.001) and every other model (all paired p < 0.003), while ESM-FP MLP drops to second-to-last (1.180 ± 0.015).

**Table 1.** Regression performance across models and splitting strategies. ElasticNet (seventh model) is omitted from main tables due to substantially inferior performance across all splits (RMSE ≥ 1.27, R^2^ ≈ 0); full ElasticNet results are reported in the project repository.

| Model | Split | RMSE (mean±SD) | R<sup>2</sup> | Pearson r | Train (s) |
| --- | --- | --- | --- | --- | --- |
| ESM-FP MLP | Random | 0.777±0.002 | 0.627±0.002 | 0.792±0.001 | 229 |
| Fusion | Random | 0.791±0.003 | 0.614±0.003 | 0.785±0.002 | 1551 |
| RF | Random | 0.819±0.000 | 0.587±0.000 | 0.769±0.000 | 87 |
| MLP | Random | 0.827±0.002 | 0.579±0.002 | 0.763±0.001 | 647 |
| GIN | Random | 0.828±0.002 | 0.578±0.002 | 0.761±0.001 | 1411 |
| XGBoost | Random | 0.894±0.001 | 0.508±0.001 | 0.717±0.001 | 113 |
| MLP | Scaffold | 0.901±0.003 | 0.498±0.003 | 0.707±0.002 | 704 |
| ESM-FP MLP | Scaffold | 0.902±0.004 | 0.497±0.005 | 0.707±0.004 | 156 |
| RF | Scaffold | 0.919±0.000 | 0.477±0.000 | 0.712±0.000 | 88 |
| Fusion | Scaffold | 0.939±0.009 | 0.455±0.010 | 0.680±0.007 | 1178 |
| GIN | Scaffold | 0.947±0.005 | 0.445±0.006 | 0.673±0.002 | 1032 |
| XGBoost | Scaffold | 0.958±0.002 | 0.433±0.002 | 0.667±0.002 | 111 |
| RF | Target | 1.066±0.002 | 0.267±0.002 | 0.521±0.002 | 76 |
| MLP | Target | 1.105±0.012 | 0.213±0.017 | 0.515±0.011 | 599 |
| XGBoost | Target | 1.131±0.004 | 0.175±0.006 | 0.432±0.006 | 97 |
| GIN | Target | 1.177±0.032 | 0.106±0.048 | 0.403±0.033 | 342 |
| ESM-FP MLP | Target | 1.180±0.015 | 0.102±0.023 | 0.388±0.020 | 44 |
| Fusion | Target | 1.196±0.031 | 0.078±0.047 | 0.392±0.034 | 683 |

**Figure 1.**
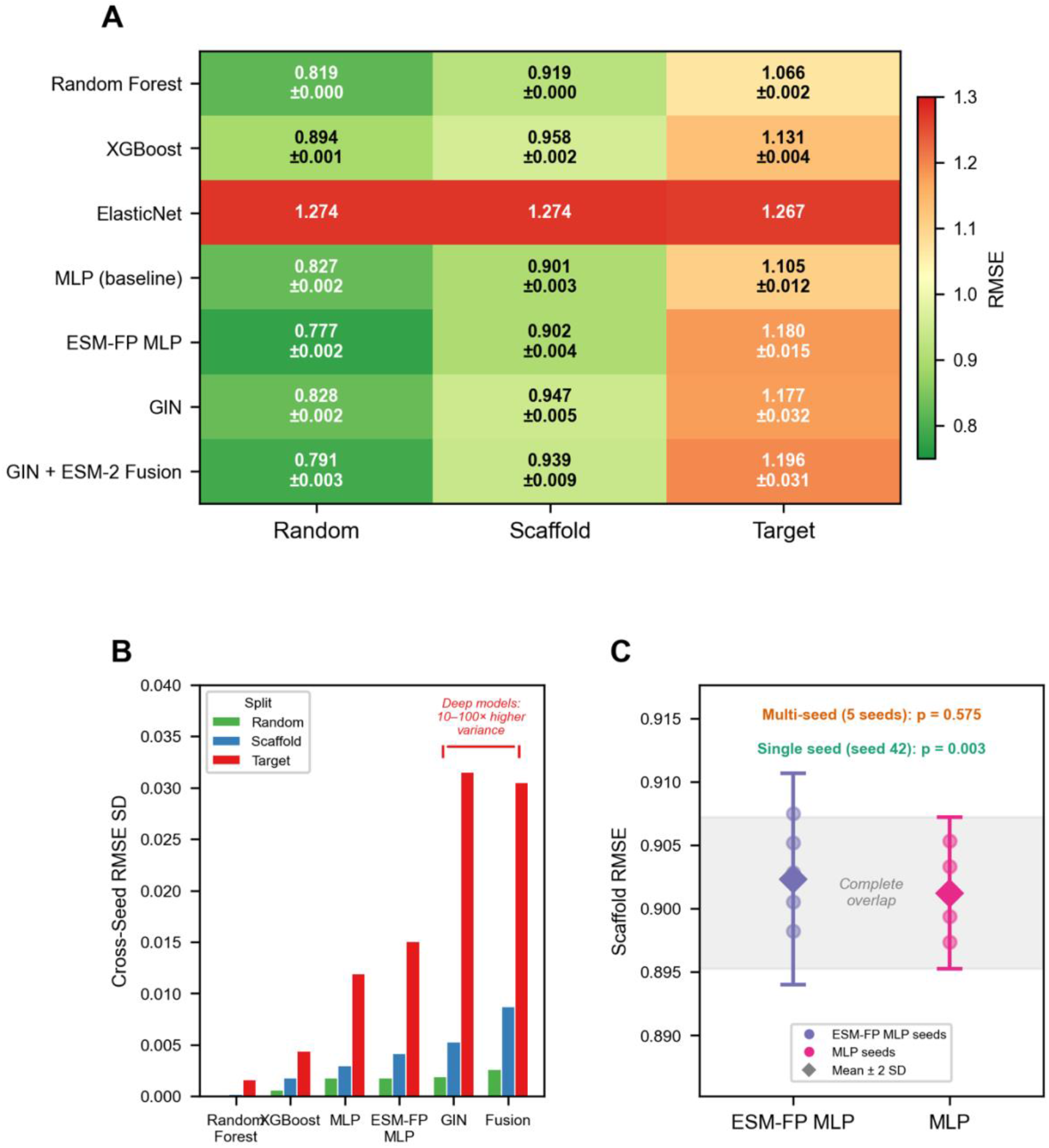
Regression metrics for the six primary comparison models across three splitting strategies. **(A)** Heatmap of test-set RMSE (multi-seed mean ± SD, 5 seeds) for all models across random, scaffold, and target-held-out splits. Color scale ranges from green (lower RMSE, better) to red (higher RMSE, worse). SD is suppressed where < 0.001. (**B**) Cross-seed RMSE standard deviation by model and split. Tree-based models (RF, XGBoost) show 10–100× lower training variance than deep models, particularly on target-held-out splits. (**C**) Per-seed scaffold RMSE for ESM-FP MLP vs. MLP (5 seeds, paired). Individual seed values (circles) and means (diamonds) show complete range overlap. Gray band indicates the overlap region.

### Performance Degradation from Random to Target Splits

**Table 2** quantifies the performance degradation using multi-seed means (**Figure 2**). ESM-FP MLP and Fusion show the largest degradation (∼51% RMSE increase), followed by GIN (42.2%). XGBoost (26.5%) and Random Forest (30.2%) exhibit the most graceful degradation. Notably, deep models show substantially higher cross-seed variance on the target split (GIN SD = 0.032, Fusion SD = 0.031) compared to tree-based models (RF SD < 0.002), meaning single-seed evaluations of deep models on hard splits can over- or under-estimate true performance by ∼3%. Early stopping behavior is consistent with this pattern. ESM-FP MLP stopped at epoch 2 on target splits versus epoch 50 on random, suggesting rapid overfitting to target-specific patterns. However, the frozen embedding strategy, limited embedding coverage (92/507 targets), and potentially insufficient regularization may also contribute.

**Table 2.** RMSE degradation from random to target-held-out splits (multi-seed mean values).

| Model | Random RMSE | Scaffold RMSE | Target RMSE | Random→Target $\Delta$ |
| --- | --- | --- | --- | --- |
| ESM-FP MLP | 0.777 | 0.902 | 1.180 | +51.8% |
| Fusion | 0.791 | 0.939 | 1.196 | +51.1% |
| GIN | 0.828 | 0.947 | 1.177 | +42.2% |
| MLP | 0.827 | 0.901 | 1.105 | +33.6% |
| Random Forest | 0.819 | 0.919 | 1.066 | +30.2% |
| XGBoost | 0.894 | 0.958 | 1.131 | +26.5% |

**Figure 2.**
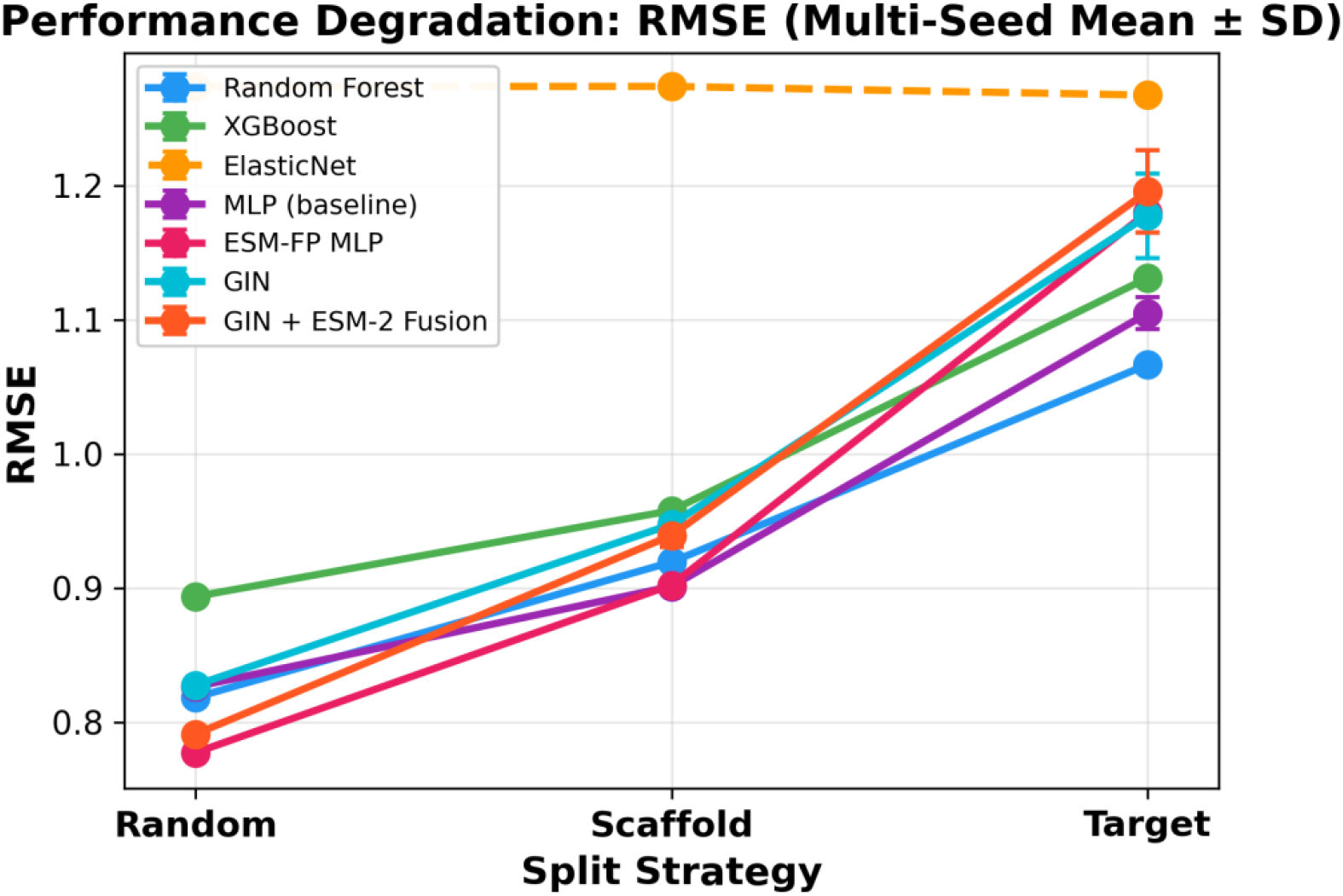
Degradation of performance for each split strategy. RMSE degradation from random to scaffold to target-held-out splits for each model (multi-seed mean ± SD, 5 seeds). Error bars represent cross-seed standard deviation. ESM-FP MLP exhibits the steepest degradation (51.8%), while XGBoost shows the lowest decline (26.5%). Models relying on protein embeddings (ESM-FP MLP, Fusion) degrade more steeply than ligand-only baselines, and also show larger error on the target split.

### Uncertainty Quantification

**Table 3** summarizes UQ results. Among the uncertainty methods evaluated here, XGBoost quantile regression showed the best calibration (miscalibration area 0.003–0.048; **Figure 3**). Random Forest tree variance showed the strongest error-uncertainty correlation (Spearman ρ = 0.334 on scaffold split) and highest selective prediction improvement (21.5% RMSE reduction at 50% retention; **Figure 4**). In our implementations, MC-Dropout yielded poorer calibration (miscalibration area 0.167–0.322), with near-zero or negative error-uncertainty correlations, suggesting that in this benchmark their uncertainty estimates provided limited practical value for selective prediction.

**Table 3.** Uncertainty quantification performance. Miscal. = miscalibration area. UQ ρ = Spearman correlation between |error| and uncertainty. Sel. Imp. = RMSE improvement at 50% retention.

| Model | Split | UQ Method | Miscal. Area | UQ $\rho$ | Sel. Imp. 50% |
| --- | --- | --- | --- | --- | --- |
| XGBoost | Random | Quantile | 0.017 | 0.195 | 12.5% |
| XGBoost | Scaffold | Quantile | 0.003 | 0.151 | 8.3% |
| XGBoost | Target | Quantile | 0.048 | 0.021 | −2.5% |
| RF | Random | Tree var. | 0.120 | 0.142 | 7.2% |
| RF | Scaffold | Tree var. | 0.037 | 0.334 | 21.5% |
| RF | Target | Tree var. | 0.166 | 0.201 | 16.3% |
| MLP | Random | Ensemble | 0.256 | 0.057 | 4.7% |
| MLP | Scaffold | Ensemble | 0.252 | 0.057 | 3.7% |
| MLP | Target | Ensemble | 0.301 | 0.126 | 7.5% |
| ESM-FP | Random | MC-Drop. | 0.167 | −0.011 | −0.5% |
| ESM-FP | Scaffold | MC-Drop. | 0.175 | −0.000 | 2.5% |
| ESM-FP | Target | MC-Drop. | 0.186 | 0.045 | 3.3% |
| GIN | Random | MC-Drop. | 0.317 | −0.030 | −0.4% |
| GIN | Scaffold | MC-Drop. | 0.322 | −0.058 | −2.1% |
| GIN | Target | MC-Drop. | 0.227 | 0.070 | 3.8% |
| Fusion | Random | MC-Drop. | 0.239 | −0.045 | −1.2% |
| Fusion | Scaffold | MC-Drop. | 0.259 | −0.061 | −2.6% |
| Fusion | Target | MC-Drop. | 0.283 | 0.042 | 5.6% |

**Figure 3.**
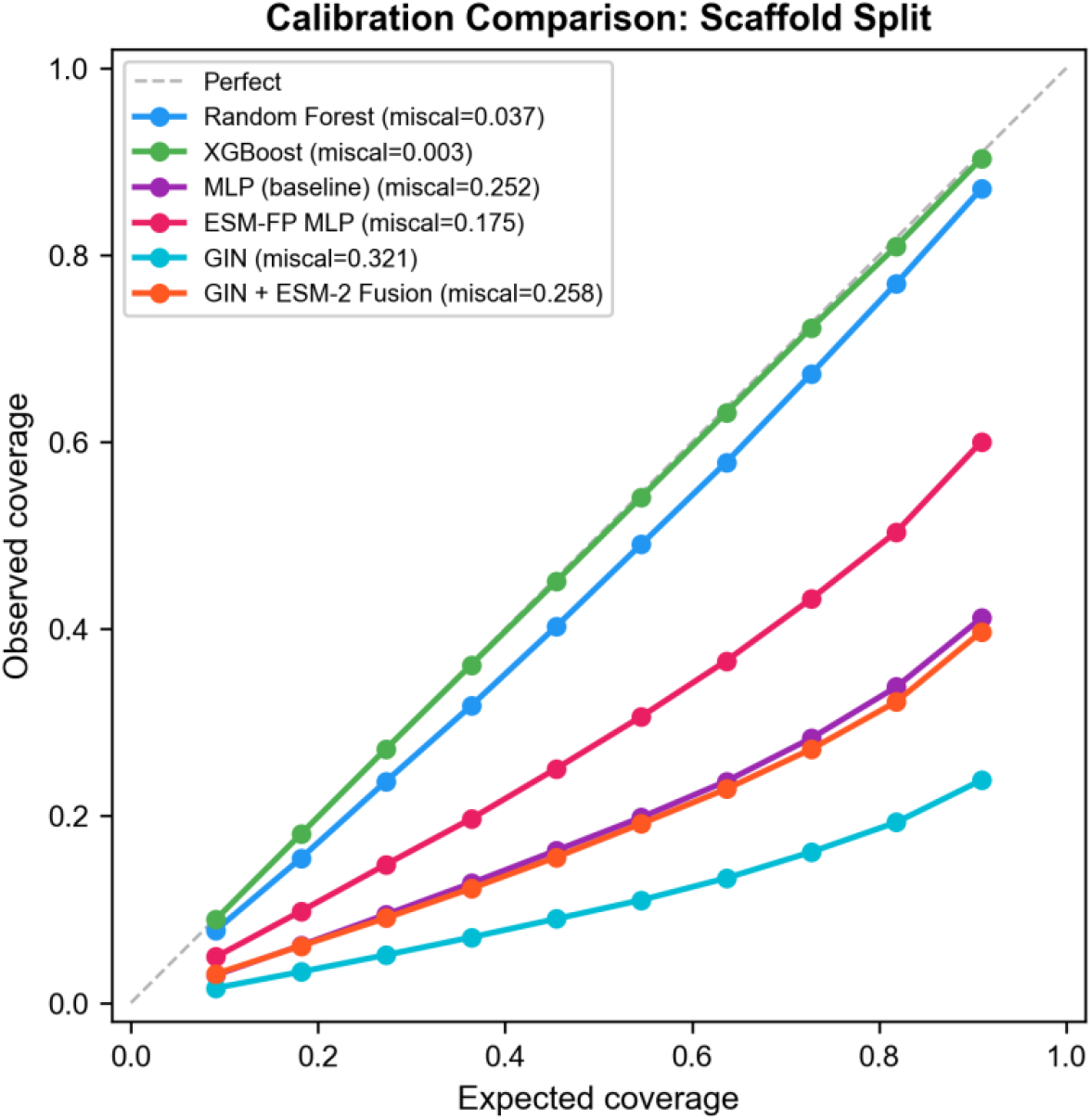
Scaffold split calibration. Calibration curves for uncertainty estimates on the scaffold split (seed 42). Each curve plots observed coverage (fraction of true values within predicted intervals) against expected coverage. XGBoost quantile regression (miscalibration area = 0.003) tracks the ideal diagonal near-perfectly. MC-Dropout from deep models (ESM-FP MLP, GIN, Fusion) falls well below the diagonal, indicating systematic overconfidence.

**Figure 4.**
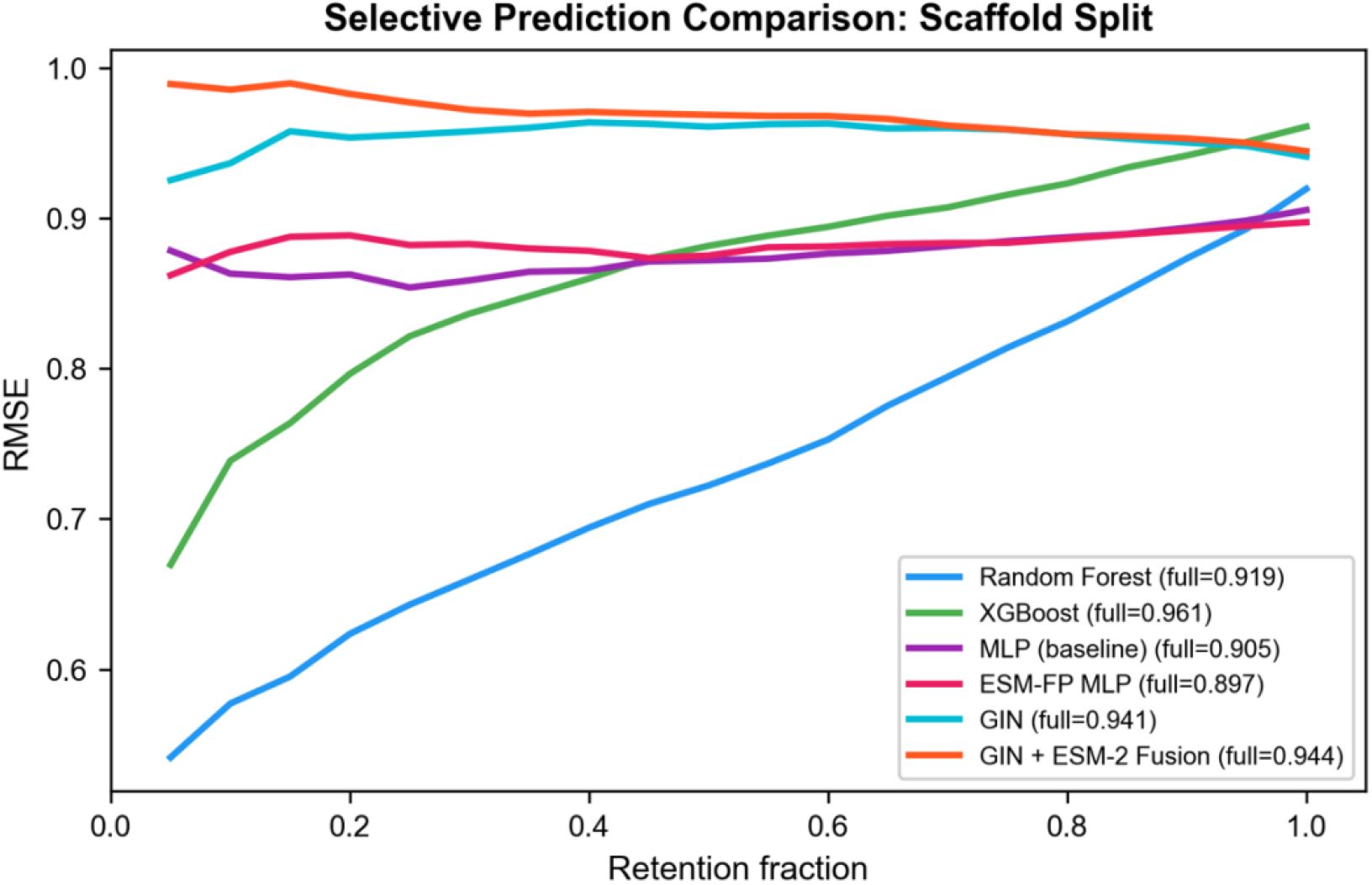
Selective prediction curves on the scaffold split (seed 42). Each curve shows test-set RMSE as a function of retention fraction, where the most uncertain predictions are removed first. Random Forest achieves the steepest improvement (21.5% RMSE reduction at 50% retention), while MC-Dropout-based models show negligible or negative benefit, confirming that their uncertainty estimates are uninformative for practical decision-making.

### JAK Family Case Study

#### Per-Target Performance

JAK targets were generally easier to predict than the full-dataset average, likely reflecting larger per-target sample sizes and denser medicinal chemistry series coverage (**Table 4, Figure 5**). ESM-FP MLP dominates on random splits, achieving RMSE = 0.566 on JAK1, 27% better than the dataset-wide average (0.777). JAK3 was consistently the hardest JAK target across all models (average RMSE 0.933 vs. 0.825 for TYK2), reflecting its broader activity range (SD = 1.35), higher apparent measurement noise (2.8% flagged), and smaller compound set (7,201 vs. 12,856 for JAK2).

**Table 4.** Per-target RMSE for JAK family members (random split). Best model per target in bold.

| Model | JAK1 (n=1053) | JAK2 (n=1216) | JAK3 (n=680) | TYK2 (n=597) |
| --- | --- | --- | --- | --- |
| ESM-FP MLP | 0.566 | 0.691 | 0.656 | 0.611 |
| Fusion | 0.625 | 0.679 | 0.738 | 0.638 |
| RF | 0.826 | 0.752 | 0.921 | 0.757 |
| MLP | 0.862 | 0.782 | 0.944 | 0.784 |
| GIN | 0.845 | 0.769 | 0.917 | 0.784 |
| XGBoost | 0.849 | 0.842 | 0.943 | 0.806 |

**Figure 5.**
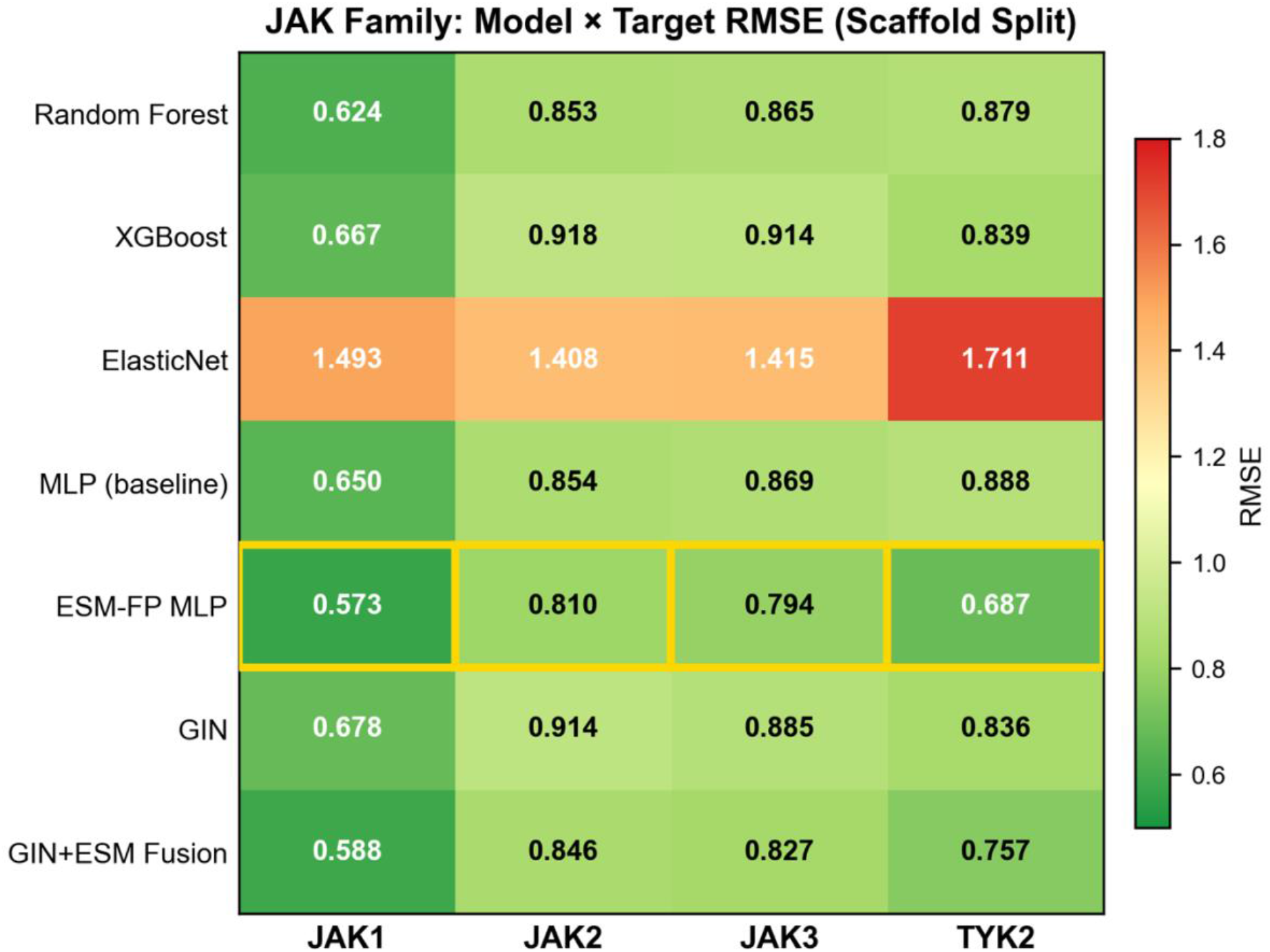
Per-target RMSE heatmap for the JAK family (scaffold split, seed 42). ESM-FP MLP and Fusion achieved the lowest RMSE for JAK1, JAK2, and TYK2, while JAK3 remains consistently the hardest target across all models.

#### Activity Cliffs

Seven activity cliff pairs were identified in the JAK test set (Tanimoto ≥ 0.85, |ΔpActivity| ≥ 1.5; **Figure 6**). The most dramatic cliff involves two stereoisomers (Tanimoto = 1.000 by 2D fingerprint) with a 2.0 log-unit JAK1 activity difference, illustrating a key limitation of the achiral fingerprint representation used here. Because Morgan fingerprints were generated without chirality encoding (includeChirality = False), these stereoisomers received identical 2048-bit vectors despite a 100-fold potency difference. JAK1 and JAK2 show more cliff pairs than JAK3 and TYK2, consistent with their larger and more densely explored compound sets. None of the top-20 worst predictions per model were flagged as noisy measurements, suggesting that these failures are not explained solely by the specific noise criterion used here.

**Figure 6.**
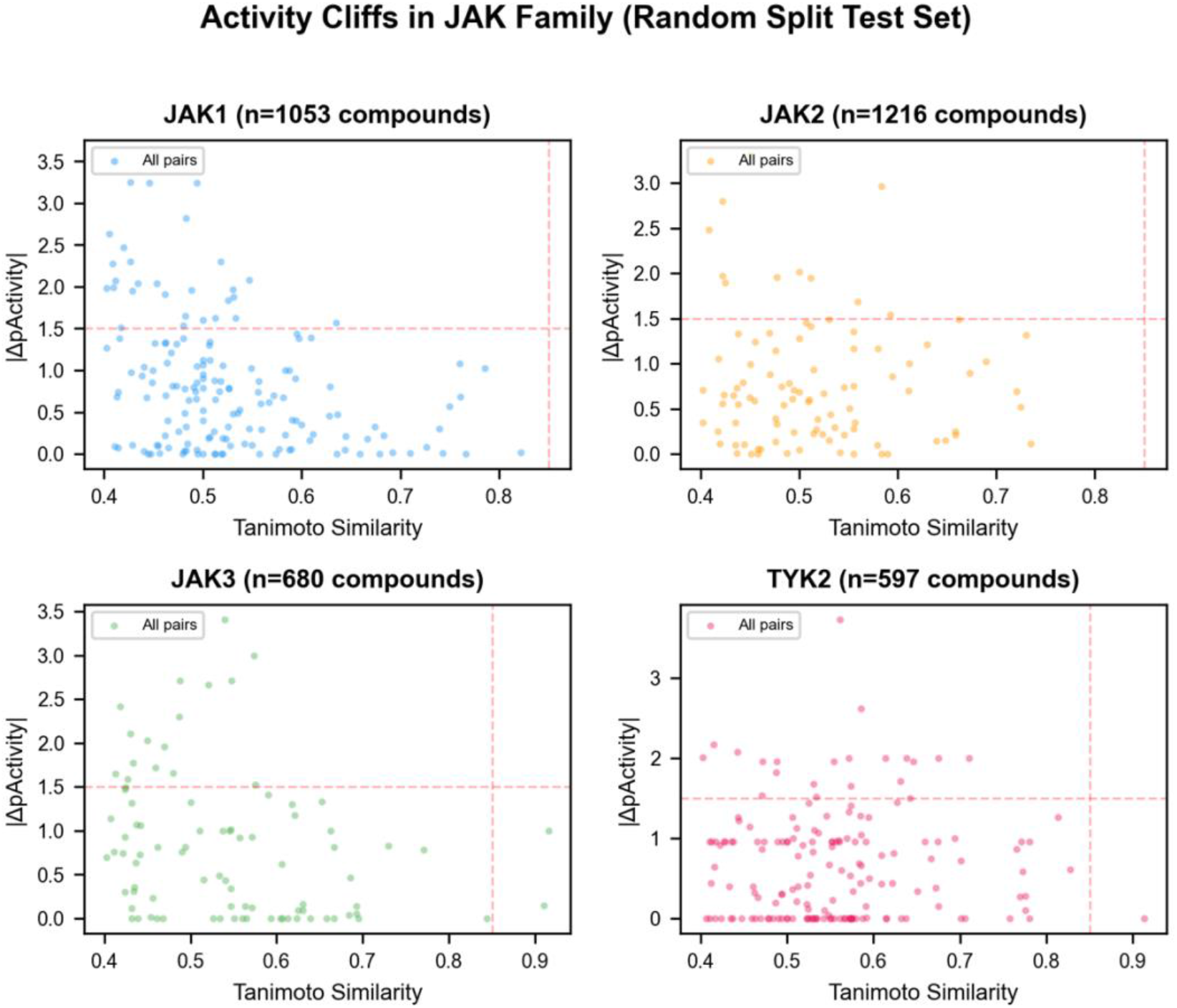
Activity cliff analysis for JAK family members (Random Split). Each panel plots |ΔpActivity| versus Tanimoto similarity for all compound pairs within a JAK target. Red points indicate activity cliffs (Tanimoto ≥ 0.85, |ΔpActivity| ≥ 1.5 log units). The most dramatic cliff is a pair of JAK1 stereoisomers (Tanimoto = 1.0) with a 2.0 log-unit potency difference, illustrating the fundamental limitation of 2D fingerprint representations.

#### Selectivity Prediction

The JAK case study also enabled a selectivity prediction analysis across family members (**Figure 7**). For 624 compounds tested against ≥ 2 JAK members under scaffold splitting, each model was asked to rank JAK members by predicted potency and identify the most potently inhibited target (**Table 5**).

**Figure 7.**
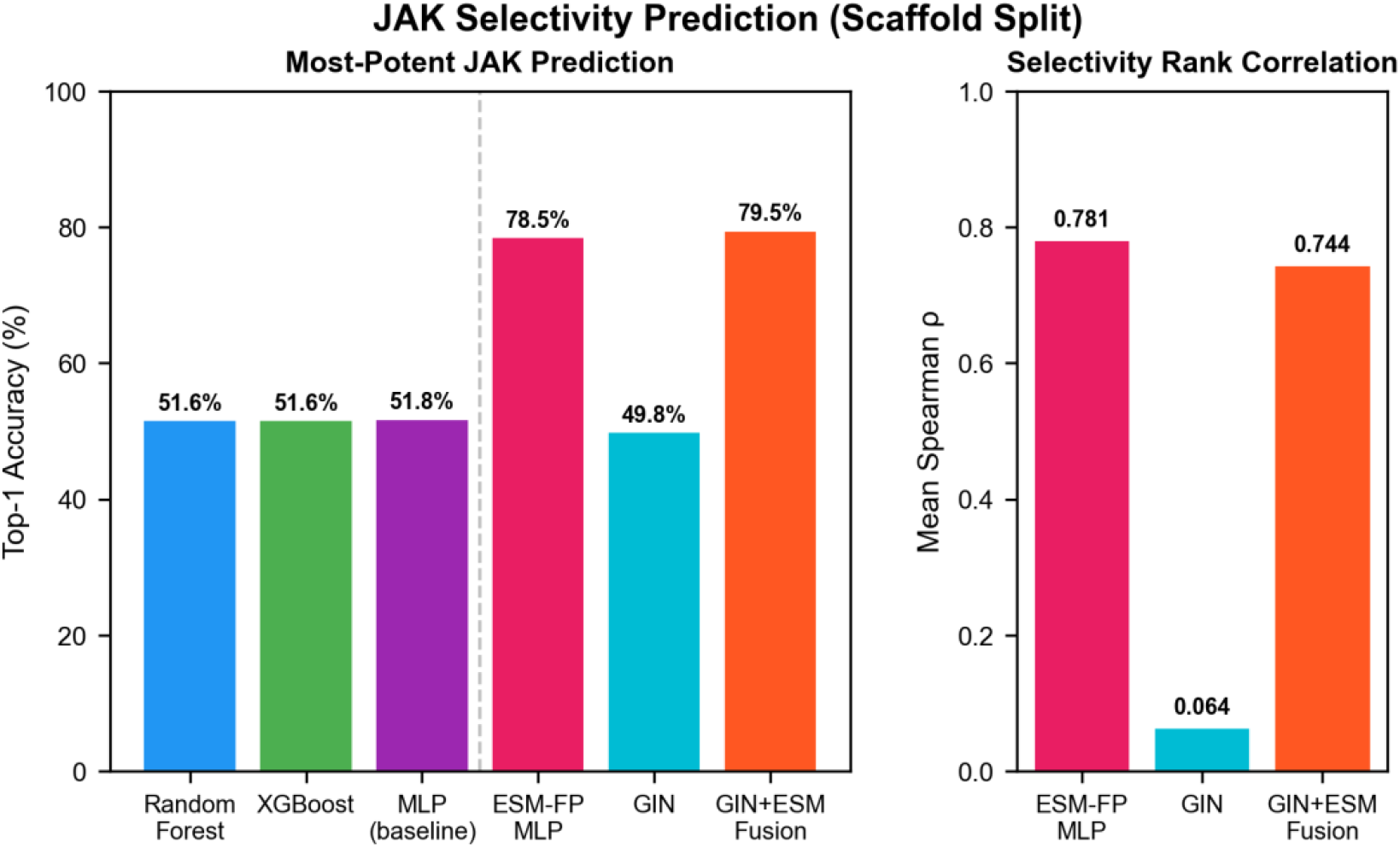
JAK selectivity prediction performance (scaffold split, seed 42, n = 624 multi-JAK compounds). Left panel: top-1 accuracy for identifying the most potently inhibited JAK member. Protein-aware models (ESM-FP MLP, Fusion) achieved ∼79% accuracy versus ∼52% for pooled fingerprint models. However, stronger baselines with explicit target identity (per-target RF, one-hot RF; Table 6) achieved 83–84%, indicating that the ESM-2 embeddings observed benefit is from implicit target identification. Right panel: Spearman rank correlation between predicted and observed selectivity profiles.

**Table 5.** JAK selectivity prediction accuracy (scaffold split, n = 624 multi-JAK compounds). Top-1 accuracy = fraction where the model correctly identifies the most potently inhibited JAK member.

| Model | Top-1 Accuracy | Mean Rank Corr. ($\rho$) | Protein Input? |
| --- | --- | --- | --- |
| Fusion | 79.5% | 0.744 | Yes |
| ESM-FP MLP | 78.5% | 0.781 | Yes |
| MLP | 51.8% | N/A | No |
| RF | 51.6% | N/A | No |
| XGBoost | 51.6% | N/A | No |
| GIN | 49.8% | 0.064 | No |

Fingerprint-based pooled models (RF, XGBoost, MLP) achieved ∼52% top-1 accuracy. These models do not directly encode target identity and produce near-identical predictions across JAK members for the same compound, making cross-target ranking effectively arbitrary. ESM-FP MLP and Fusion achieved ∼79% accuracy by incorporating ESM-2 protein embeddings that encode the sequence differences between JAK family members. GIN (49.8%) also fails because, despite operating on molecular graphs, it lacks protein input. These results persist under scaffold splitting, suggesting that it is not explained solely by close analogue leakage.

A concern with this comparison is that the pooled fingerprint models were not designed for selectivity as they lack any target-specific input. To provide a more relevant comparison, we implemented stronger selectivity baselines that give fingerprint models explicit target identity through three strategies: separate per-target models, one-hot target encoding appended to the fingerprint, and pairwise ΔpActivity difference models.

### ESM-2 Embedding Coverage: Ablation Studies

A major limitation of the protein-aware models is that only 92 of 507 kinase targets (18.1%) have real ESM-2 embeddings; the remaining 415 targets received a fallback embedding (row 0 of the embedding matrix—an arbitrary kinase’s representation). To quantify the impact, we conducted two ablation studies.

#### Fallback strategy ablation

We retrained ESM-FP MLP and Fusion with three fallback strategies for the 415 uncovered targets: row0 (original arbitrary embedding), zero vector (no protein signal), and mean of all 92 real embeddings (“generic kinase”). For 5 of 6 model×split conditions, the fallback strategy had minimal impact (Δ < 0.02 RMSE), confirming that the models learned to effectively ignore the protein branch for fallback targets. The one exception is Fusion on the target split, where the mean-vector fallback caused substantial degradation (RMSE = 1.239 vs. 1.115 for zero), suggesting sensitivity to misleading protein signals when predicting novel kinase subfamilies.

#### Clean 92-target subset

To isolate the effect of real protein embeddings from fallback noise, we retrained all models on a clean subset containing only the 92 targets with real ESM-2 embeddings (67,902 records, 19.2% of the full dataset). The results were notable, ESM-FP MLP achieves RMSE = 0.647 on random splits, a 16.7% improvement over the full-dataset result (0.777), and 0.826 on scaffold splits (vs. 0.902). Fusion similarly improves from 0.791 to 0.695 for random-split and from 1.196 to 1.062 target-split. Baseline models also improve on the smaller subset (RF: 0.819 to 0.786), likely because the 92 well-studied targets have cleaner data. This suggests that the 92-target results should be interpreted as an upper bound on protein embedding value. Notably, ESM-FP MLP degrades on the 92-target target split (1.180 to 1.260), possibly because with only about 10 held-out targets the test set is too small for reliable evaluation.

These ablation results suggest that incomplete ESM-2 coverage likely masked some of the potential value of protein-aware models in the full-dataset analysis. However, because the 92-target subset is enriched for well-studied kinases with denser SAR coverage, these results should be interpreted as an upper-bound estimate of protein embedding value, not a direct like-for-like replacement of the full-dataset benchmark. Expanding UniProt coverage to the remaining 415 targets via NCBI Protein, PDB, or manual curation remains the single highest-priority improvement for protein-aware architectures. The JAK case study is not directly affected by the fallback limitation as all four JAK kinases have distinct ESM-2 embeddings.

## Discussion

### Evaluation Methodology and Reported Performance

In this benchmark, evaluation methodology had as much or more impact on reported performance than the choice of model architecture. The best model on random splits (ESM-FP MLP, RMSE = 0.777 ± 0.002 across 5 seeds) degrades to RMSE = 1.180 ± 0.015 on target splits. Model rankings reverse: ESM-FP MLP ranks first on random splits but second-to-last on target splits, while Random Forest ranks third on random but first on target; both multi-seed paired t-tests (all p < 0.003) and bootstrap win-rate matrices confirm these rankings are stable. Critically, the apparent ESM-FP MLP advantage over MLP on scaffold splits—initially reported as significant by single-seed bootstrap (p = 0.003)—was revealed as a false positive by multi-seed evaluation (p = 0.575), demonstrating that single-seed comparisons of deep models can be misleading, particularly on harder splits. The 12–52% increase in RMSE from random to target-held-out splits demonstrates how strongly reported performance depends on whether close chemical and target-family relationships are preserved across train and test partitions. These findings are consistent with Luo et al.^16^, who showed that deep learning models for kinase bioactivity prediction exhibit limited generalization when information leakage is controlled, and extend that work by demonstrating the phenomenon across a broader range of architectures including protein-aware models.

### Protein Embeddings: Interpolation, Extrapolation, and Selectivity

ESM-2 protein embeddings present a nuanced picture across three use cases. For interpolation (random splits, known targets), they improve RMSE by 5.1% over RF (0.777 ± 0.002 vs. 0.819 ± 0.000; paired t-test p < 0.001). For extrapolation to held-out kinase subfamilies, the frozen ESM-based models underperformed the ligand-only baselines evaluated here, ESM-FP MLP drops to second-to-last place with 51.8% RMSE degradation (1.180 ± 0.015), among the worst of any model (paired p < 0.001 vs. RF). Early stopping at epoch 2 on target splits is consistent with poor transfer, although other factors, including limited embedding coverage (92/507 targets), the frozen embedding strategy, and potentially insufficient regularization may also contribute. We note that this result does not demonstrate that protein embeddings are inherently harmful for extrapolation. Instead in this specific benchmark configuration, frozen ESM-2 and shared arbitrary-kinase fallback for 82% of targets failed to improve generalization.

The JAK case study initially appeared to reveal a third use case where protein embeddings provide their clearest advantage for selectivity prediction. ESM-FP MLP and Fusion achieved 79% top-1 accuracy versus 52% for pooled fingerprint models at identifying the most potently inhibited JAK member.

However, stronger baselines with explicit target identity per-target models (83.0%), one-hot target encoding (83.5%), and pairwise difference models (82.7%) match or exceed the protein-aware models (**Table 6**). This demonstrates that the ESM-2 embedding functions primarily as an implicit target identifier. In this context embeddings provide the model with target-discriminative information, but simpler one-hot encoding achieves the same result more effectively.

**Table 6.** Stronger selectivity baselines with explicit target identity. Scaffold split, n = 624 multi-JAK compounds.

| Model | Top-1 Accuracy | Rank Corr. ($\rho$) | Target Info |
| --- | --- | --- | --- |
| Per-target RF | 83.0% | 0.833 | Separate models |
| One-hot RF (FP + target) | 83.5% | 0.813 | One-hot encoding |
| One-hot MLP (FP + target) | 82.9% | 0.825 | One-hot encoding |
| Pairwise RF ($\Delta p$Activity) | 82.7% | 0.791 | Pairwise differences |
| Per-target MLP | 75.2% | 0.695 | Separate models |
| ESM-FP MLP | 78.5% | 0.781 | ESM-2 embedding |
| Fusion | 79.5% | 0.744 | ESM-2 + GIN |

The true value proposition of protein embeddings for selectivity would emerge in a project where the model must generalize to targets not seen during training where one-hot encoding is unavailable by definition. However, this is the setting where frozen ESM-2 models underperform in this benchmark. Resolving this conflict between protein embeddings that transfer to novel targets remains an ongoing problem.

These results together suggest that frozen ESM-2 embeddings encode useful relative information about target identity within a protein family but, at least in this study, do not capture the physicochemical context of binding that would transfer to structurally distant kinases. The ESM-2 ablation studies suggest that incomplete embedding coverage (92/507 targets) likely masked some of the potential value of protein-aware models. On the clean 92-target subset, ESM-FP MLP achieves RMSE = 0.647, a 16.7% improvement over the full-dataset result, though this subset is enriched for heavily studied targets and should be understood as an upper bound. Recent work by Prabakaran and Bromberg^9^ demonstrates that protein language model embeddings vary substantially in quality across the proteome, their Random Neighbor Score (RNS) metric reveals that 19–46% of human proteins are underlearned by current pLMs, with embeddings for these proteins indistinguishable from random sequences in latent space. This finding provides independent mechanistic support for our observation that incomplete ESM-2 coverage degrades downstream prediction. When 82% of targets receive a shared fallback vector, the model effectively loses access to protein-level information for the majority of the dataset. RNS-based screening could further refine our ablation analysis by stratifying even the 92 covered targets by embedding confidence. Fine-tuning the last few ESM-2 layers, expanding embedding coverage to all 507 targets, or incorporating 3D binding-site structural information may be necessary to realize the extrapolation potential of protein-aware models.

### Learned vs. Fixed Molecular Representations

GIN did not outperform the Morgan fingerprint Random Forest baseline on any split in this benchmark (GIN random RMSE = 0.828 ± 0.002 vs. RF = 0.819 ± 0.000; paired t-test p < 0.001 favoring RF). MLP (0.827 ± 0.002) and GIN are not statistically different on random splits (p = 0.403), but MLP significantly outperforms GIN on scaffold splits (ΔRMSE = −0.046, p < 0.001) and target splits (ΔRMSE = −0.073, p = 0.013). At ∼350K records, the 3-layer GINConv architecture with 128-dim hidden states does not appear to learn representations that meaningfully exceed what 2,048-bit ECFP4 fingerprints already capture, though we acknowledge that deeper or attention-based graph architectures might perform differently. The JAK case study reinforces this, GIN also fails at selectivity prediction (49.8% accuracy), confirming that the ESM-FP MLP/Fusion advantage is a protein-representation advantage, not a ligand-representation advantage.

### Uncertainty Quantification

The UQ results in this benchmark suggest a practical hierarchy, among the uncertainty methods evaluated here, XGBoost quantile regression showed the best calibration (near-perfect on scaffold splits), Random Forest tree variance provided the most actionable uncertainty for selective prediction (21.5% RMSE reduction at 50% retention), and in our implementations, MC-Dropout yielded poorer calibration and weaker selective prediction utility than the tree-based uncertainty baselines. The finding that MC-Dropout produces negative error-uncertainty correlations in several settings, meaning more uncertain predictions are not actually less accurate, suggests that in this benchmark their uncertainty estimates provided limited practical value for selective prediction^17^. For practical use, Random Forest’s combination of competitive point predictions and well-correlated uncertainty makes it the most operationally useful model evaluated here.

### Implications for Drug Discovery ML Benchmarking

Our results support several practical recommendations. First, benchmark evaluations should always report scaffold and target-based splits alongside random splits. Second, simple baselines like Random Forest with Morgan fingerprints, should be mandatory comparisons. Third, multi-seed evaluation (≥3 seeds with mean ± SD reporting) is essential for deep model comparisons, our scaffold-split false positive demonstrates that single-seed results can mislead. Fourth, uncertainty quantification should be evaluated alongside point predictions. Fifth, protein-aware architectures should be evaluated specifically on target-held-out splits and on selectivity tasks with strong baselines as these represent distinct use cases that may yield different conclusions about the value of protein information.

### Limitations

Because the benchmark pools IC_50_, K_i_, and K_D_ values into a common pActivity target, it should be interpreted as a mixed-endpoint bioactivity benchmark with endpoint-dependent heterogeneity. Restricting to a single endpoint type or adding assay-type variables would improve interpretability. ESM-2 embeddings were available for only 92 of 507 kinase targets (18.1%). The remaining targets received a shared fallback embedding, an arbitrary kinase’s real ESM-2 vector reused as a placeholder, meaning protein-aware models operated with non-target-specific protein information for the majority of targets. Our ablation studies suggest that incomplete coverage likely masked some of the potential value of protein-aware models, ESM-FP MLP achieves 16.7% lower RMSE on the clean 92-target subset, but because those 92 targets are likely enriched for well-studied kinases with cleaner SAR trends, this result should be interpreted as an upper-bound estimate rather than a direct replacement of the full-dataset benchmark. Expanding UniProt coverage remains the highest-priority improvement for protein-aware architectures. Additionally, applying embedding quality metrics such as the Random Neighbor Score to screen or weight embeddings by confidence could improve reliability even for targets with nominal coverage. We did not evaluate temporal splits^10^, which are an important test of deployment realism in medicinal chemistry settings. Our GIN architecture is relatively shallow, deeper or attention-based graph architectures may perform better. Multi-seed evaluation addresses model-training variability, the primary source of uncertainty for deep models, but does not capture partition sensitivity. All seeds share the same scaffold assignment and target-family split. Because scaffold and target partitions are determined by a single random seed (seed 42), model rankings under alternative partitions could differ, and this remains the principal methodological limitation not yet addressed. The JAK case study represents a single well-studied subfamily, generalization to less-characterized kinase families remains to be established.

Selectivity prediction was evaluated only within the JAK family and requires broader validation, in particular, evaluation on targets not seen during training where protein embeddings advantage over one-hot encoding would be most meaningful.

## Conclusions

Through systematic evaluation of seven ML architectures across three splitting strategies on kinase bioactivity records, with multi-seed replication, bootstrap confidence intervals, and paired significance tests, we show that evaluation methodology is a major determinant of reported benchmark performance in kinase bioactivity prediction. In this benchmark, complex protein-aware models achieved significant improvements under lenient evaluation but fail to outperform simple baselines under the most stringent conditions. Notably, the single-seed scaffold-split advantage of ESM-FP MLP over MLP was revealed as a false positive by multi-seed evaluation, underscoring the necessity of replicated training for deep model comparisons. Random Forest with Morgan fingerprints emerged as the most robust and practically useful baseline for absolute pActivity prediction, offering competitive accuracy, superior uncertainty quantification, near-zero training variance, and training times under 2 minutes.

For selectivity prediction, ESM-FP MLP and Fusion achieved 79% top-1 accuracy at identifying the most potently inhibited JAK family member, versus 52% for pooled fingerprint models lacking target-specific input. However, stronger baselines with explicit target identity per-target models and one-hot target-conditioned models, match or exceed the protein-aware models, demonstrating that the ESM-2 embedding functions primarily as an implicit target identifier in this context. The ablation studies support this conclusion, restricting to targets with genuine ESM-2 coverage substantially improves protein-aware model performance, the enrichment of well-studied kinases in this subset means the 16.7% improvement should be treated as an upper bound. Expanding embedding coverage and developing protein-aware architectures that generalize to unseen targets remain the key open challenges for the field.

### Practical recommendation

For standard kinase pActivity prediction, Random Forest or MLP with Morgan fingerprints provides robust, well-calibrated predictions with minimal computational overhead and near-zero training variance. Protein-aware models should be reserved for settings where target identity must be explicitly encoded, with the caveat that simpler target-conditioned approaches may perform equally well when the targets of interest are known. Any comparison involving deep models should be supported by multi-seed evaluation (≥3 seeds), as single-seed point estimates can produce false positives sufficient to change published conclusions.

## Data and Code Availability

The complete codebase, including data curation pipelines, feature engineering, model training, evaluation, and visualization scripts, is available at https://github.com/jmabbott40/kinase-affinity-baselines. Raw bioactivity data were sourced from ChEMBL (public domain). Pre-computed features and trained model weights are deposited at 10.5281/zenodo.19614454. All experiments are reproducible using the provided configuration files and random seeds.

## Acknowledgments

Analysis was performed on an AWS g5.12xlarge instance using the Deep Learning OSS Nvidia Driver AMI GPU PyTorch 2.7 (Ubuntu 22.04) 20260118. Claude Code was utilized to assist with building analysis pipeline. Activity data was obtained from ChEMBL 36. This work was conducted independently of authors current employer and does not represent the views of any institution or employer.

## Appendix A: Per-Seed Regression Results

**Table S1** reports the range (min–max) and spread of RMSE and R^2^ across 5 training seeds for all models and splits. Full per-seed values are available in the project repository (results/supplement_tables/S6_multi_seed_detailed.csv). Deep models exhibit substantially higher cross-seed variance than tree-based models, particularly on the target split (GIN SD = 0.032 vs. RF SD = 0.002). The ESM-FP MLP vs. MLP scaffold-split comparison is highlighted: their RMSE ranges overlap completely (ESM-FP MLP: 0.898–0.908; MLP: 0.897–0.905), confirming that the single-seed false positive (p = 0.003) was driven by a particular initialization rather than a systematic advantage.

**Table S1.** Multi-seed RMSE summary (5 seeds: 42, 123, 456, 789, 1024). SD = standard deviation across seeds.

| Model | Split | RMSE Mean | SD | Min | Max | Range |
| --- | --- | --- | --- | --- | --- | --- |
| ESM-FP MLP | Random | 0.777 | 0.002 | 0.775 | 0.780 | 0.005 |
| Fusion | Random | 0.791 | 0.003 | 0.788 | 0.795 | 0.007 |
| RF | Random | 0.819 | 0.000 | 0.818 | 0.819 | 0.001 |
| MLP | Random | 0.827 | 0.002 | 0.824 | 0.829 | 0.005 |
| GIN | Random | 0.828 | 0.002 | 0.826 | 0.831 | 0.005 |
| XGBoost | Random | 0.894 | 0.001 | 0.893 | 0.895 | 0.002 |
| MLP | Scaffold | 0.901 | 0.003 | 0.897 | 0.905 | 0.008 |
| ESM-FP MLP | Scaffold | 0.902 | 0.004 | 0.898 | 0.908 | 0.010 |
| RF | Scaffold | 0.919 | 0.000 | 0.919 | 0.920 | 0.001 |
| Fusion | Scaffold | 0.939 | 0.009 | 0.930 | 0.949 | 0.019 |
| GIN | Scaffold | 0.947 | 0.005 | 0.940 | 0.954 | 0.014 |
| XGBoost | Scaffold | 0.958 | 0.002 | 0.956 | 0.961 | 0.005 |
| RF | Target | 1.066 | 0.002 | 1.064 | 1.068 | 0.004 |
| MLP | Target | 1.105 | 0.012 | 1.090 | 1.119 | 0.029 |
| XGBoost | Target | 1.131 | 0.004 | 1.124 | 1.135 | 0.011 |
| GIN | Target | 1.177 | 0.032 | 1.139 | 1.222 | 0.083 |
| ESM-FP MLP | Target | 1.180 | 0.015 | 1.164 | 1.199 | 0.035 |
| Fusion | Target | 1.196 | 0.031 | 1.149 | 1.231 | 0.082 |

